# Human inbreeding has decreased in time through the Holocene

**DOI:** 10.1101/2020.09.24.311597

**Authors:** Francisco C. Ceballos, Kanat Gürün, N. Ezgi Altınışık, Hasan Can Gemici, Cansu Karamurat, Dilek Koptekin, Kıvılcım Başak Vural, Elif Sürer, Yılmaz Selim Erdal, Anders Götherström, Füsun Özer, Çiğdem Atakuman, Mehmet Somel

## Abstract

The history of human inbreeding is controversial. The development of sedentary agricultural societies may have had opposite influences on inbreeding levels. On the one hand, agriculture and food surplus may have diminished inbreeding by increasing population sizes and lowering endogamy, i.e. inbreeding due to population isolation. On the other hand, increased sedentism, as well as the advent of private property may have promoted inbreeding through the emergence of consanguineous marriage customs or via ethnic and caste endogamy. The net impact is unknown, and to date, no systematic study on the temporal frequency of inbreeding in human societies has been conducted. Here we present a new approach for reliable estimation of runs of homozygosity (ROH) in genomes with ≥3x mean coverage across >1 million SNPs, and apply this to 440 ancient Eurasian genomes from the last 15,000 years. We show that the frequency of inbreeding, as measured by ROH, has decreased over time. The strongest effect is associated with the Neolithic transition, but the trend has since continued, indicating a population size effect on inbreeding prevalence. We further show that most inbreeding in our historical sample can be attributed to endogamy, although singular cases of high consanguinity can also be found in the archaeogenomic record.

**Highlights:** A study of 440 ancient genomes shows inbreeding decreased over time.

The decrease appears linked with population size increase due to farming.

Extreme consanguineous matings did occur among farmers, but rarely.

## RESULTS AND DISCUSSION

To study ROH levels in time, we tailored the *PLINK* implementation of ROH calling to suit low coverage ancient genomes. Simulations performed by downsampling 44 relatively high coverage (>10x) ancient genomes revealed that the default *PLINK* algorithm overestimates the sum and number of ROH’s at <4x coverage, due to missed heterozygous positions in the data (Table S1, Figure S1, Figure S2 A and C). We accounted for this effect using an empirical approach: depending on coverage, we vary the parameters of *PLINK* with respect to the number of heterozygous SNP allowed per window (see Materials and Methods). With this approach we were able to estimate the number and sum of ROH >1Mb reliably for >3x coverage genomes in simulations (Figure S1 B and D). We further confirmed that ROH calls >1Mb were free of the influence of coverage by studying the correlation between genome coverage and the sum or the number of ROH >1Mb across 440 ancient individuals with ≥3x mean SNP coverage across the 1240K SNP set [1] (see Materials and Methods). Meanwhile, we discarded small ROH (<1Mb), as we found that these cannot be identified reliably with low coverage genomes using our approach (see Materials and Methods).

Our analyses below focus on the number and sum of ROH >1Mb estimated across these n=440 published genomes using our empirical approach (Figure 1, Figure 2A), as well as among n=444 contemporary human individuals (Figure 2B). We focused on West Eurasia (Europe) and Central Eurasia (SW Asia, Caucasus, and Central Asia), regions with the highest published ancient genome data density. To study the effects of changing sociocultural organization through time, we separated past societies into four historical categories based on the degree of their social complexity: hunter-gatherers, who subsisted on the wild resources of the land within egalitarian mobile bands (e.g. Gravettian hunter-gatherers in Eastern Europe); simple agriculturalists, the earliest adopters of agriculture within relatively egalitarian sedentary communities (e.g. Linearbandkeramik farmers of Central Europe); early complex agriculturalists, farmer/pastoralist communities with an emerging institutionalized hierarchy and specialization (e.g. Bell Beaker groups known mainly from burials in West and Central Europe); and advanced complex agriculturalists, who lived in highly stratified societies organized around state systems (e.g. the Roman state in the Mediterranean).

**Figure 1.**
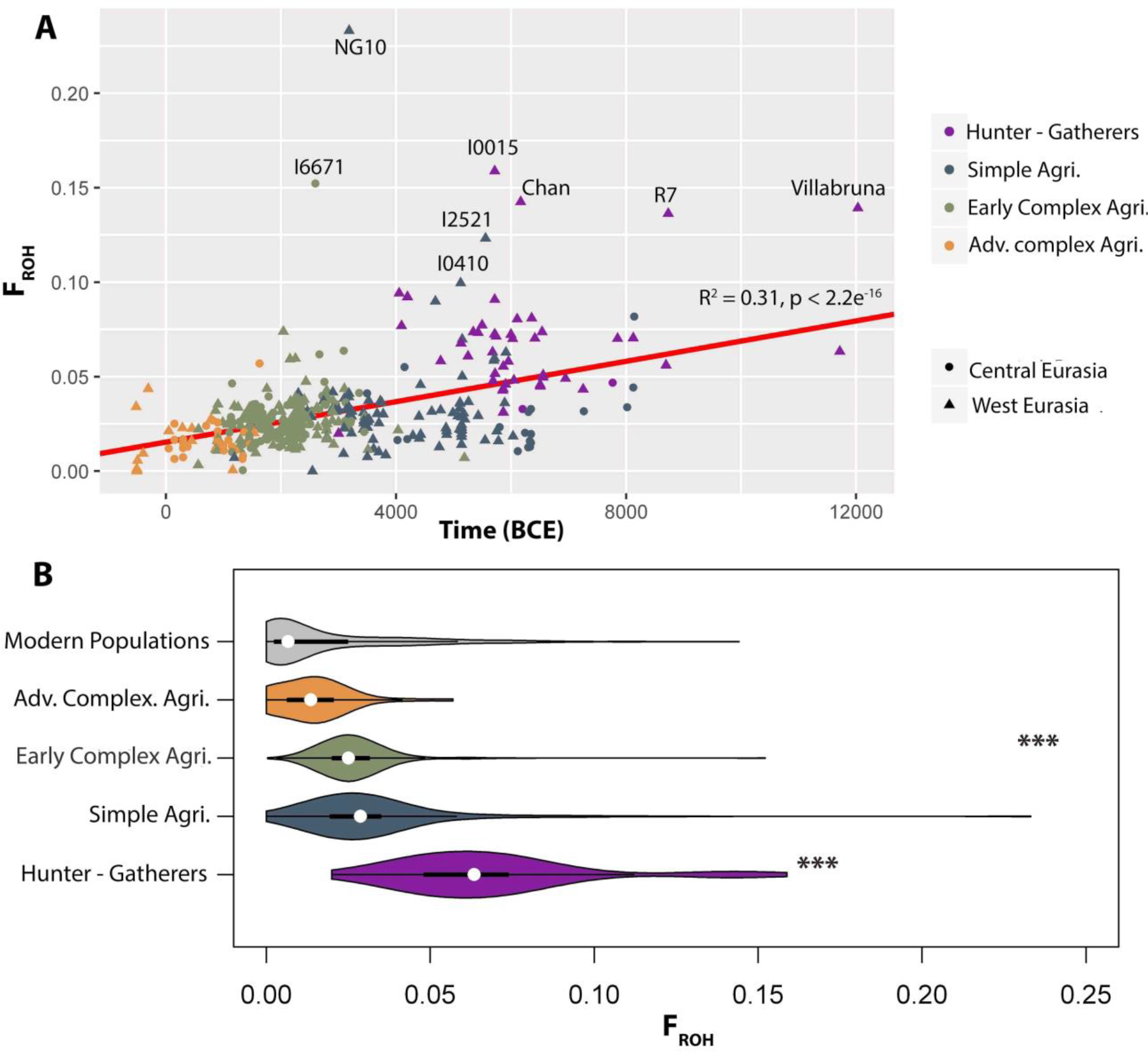
Temporal distribution of the genomic inbreeding coefficient (F_ROH_). (**A**) Regression of F_*ROH*_ estimates against time in years Before the Common Era (BCE). Historical categories are defined with colours: Hunter-gatherers in violet, simple agriculturalists (Simple Agri.) in blue, early complex agriculturalists (Early Complex Agri.) in green, advanced complex agriculturalists (Adv. Complex Agri.) in orange, present-day populations from the Human Genome Diversity Panel in grey. Region of origin of each individual is shown with a symbol: Central Eurasia with a circle and West Eurasia with a triangle. The regression line was obtained by analysing only the ancient individuals (n=440) and has a significant slope (*β*_Time_ = 6.09e^−6^, *p* = 2e^−16^, *R*^2^ = 0.31, *p* < 2.2e^−16^). (**B**) Violin plots of F_*ROH*_ estimates for the different historical categories and present-day populations from the Human Genome diversity Panel. Asterisks represent significance (<0.001) calculated by the pairwise Wilcoxon rank sum test with continuity correction. We detected a significant difference between hunter-gatherers and the rest of the groups, and between early complex agriculturalists and advanced complex agriculturalists or HGDP’s modern-day populations.

**Figure 2.**
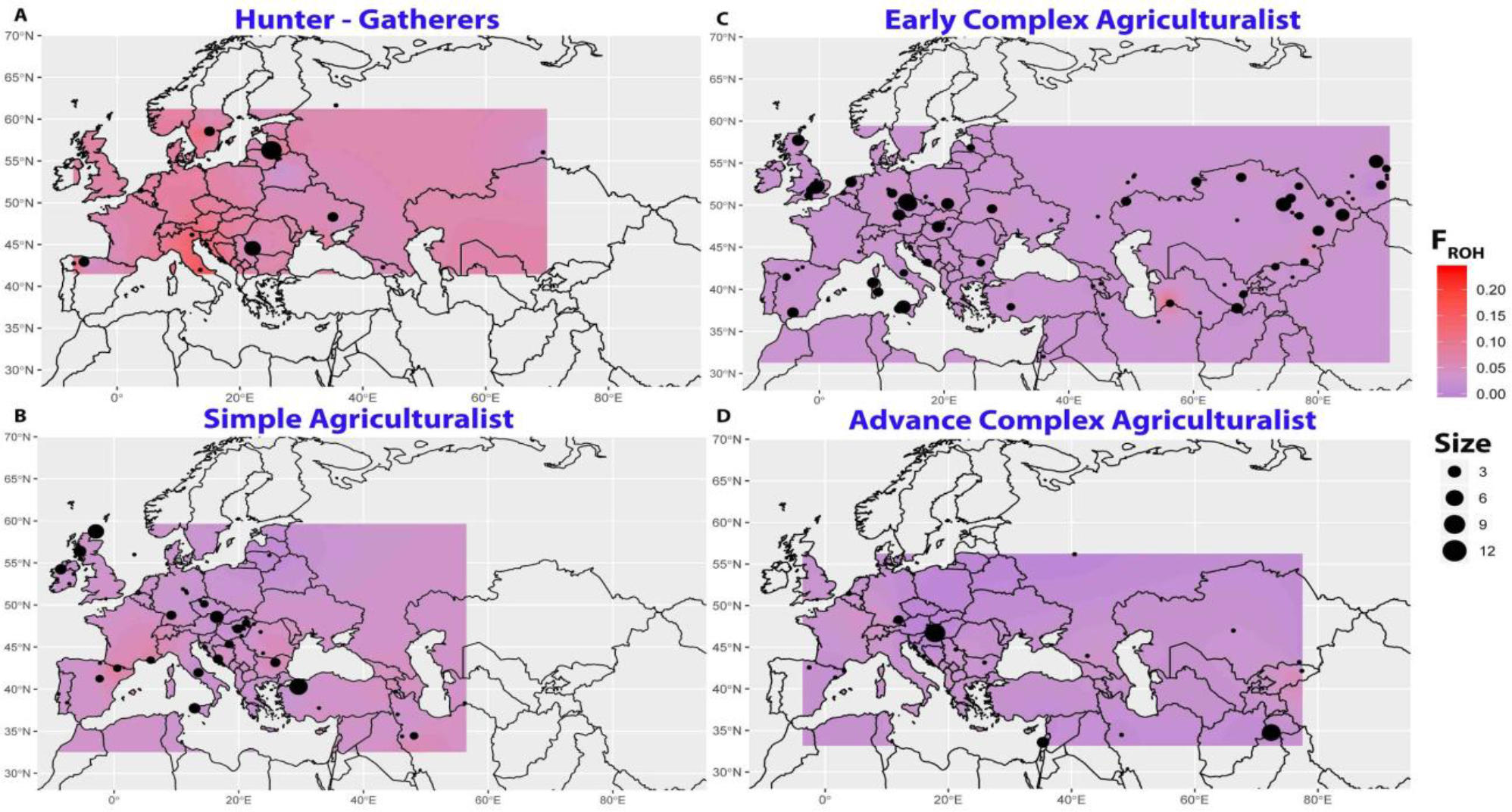
Spatially kriged reconstructions for the distribution of the genomic inbreeding coefficient (F_*ROH*_). The colors represent the predicted F_*ROH*_ values. The panels show spatial kriging of F_*ROH*_ estimates in hunter-gatherers (A), in the simple agriculturalists (B), in the early complex agriculturalists (C), and in the advanced complex agriculturalists (D).

### Temporal and spatial distribution of human inbreeding

We first studied the temporal distribution of *F*_*ROH*_, or genomic inbreeding levels. We find a manifest trend of decreasing levels of inbreeding over time in West and Central Eurasia (Figure 1A). When separating the data into five historical categories, from hunter-gatherers to advanced complex agriculturalists and finally to contemporary humans, we observe the same trend. Notably, the largest shift in *F*_*ROH*_ occurs between hunter-gatherers and agriculturalists, during the Neolithic transition, but the trend is sustained in later periods. We find significant differences between almost every pair of historical categories but simple agriculturalists vs. early complex agriculturalists, and advanced complex agriculturalists vs. modern-day populations (Table S2).

We further analysed the data using a multiple regression model with *F*_*ROH*_ as the dependent variable, and time (i.e. historical age) and historical category as independent variables. Time had a positive and significant effect (*β*_Time_ = 3.98e^−06^, *p* = 3.7e^−06^), while the effect of the different historical categories was also significant in comparison to the baseline set by the hunter-gatherers (simple agriculturalists’ effect = −3.01e^−02^, *p* < 2.2e^−16^; early complex agriculturalists’ effect = −2.5e^−02^, *p* = 7.1e^−08^; advanced complex agriculturalists’ effect = −3.1e^−02^, *p* = 5.3e^−07^ and contemporary societies’ effect = −3.4e^−02^, *p* = 1.83e^−05^). This can also be observed in Figure 1B. Notably, contemporary populations have the lowest inbreeding levels, despite notable variability within this group (Table 1).

**Table 1.** Summary statistics for the genomic inbreeding coefficient calculated from ROH (*F*_*ROH*_) across historical categories and geographical regions. N: number of individuals. IQC: interquartile range. F>0.0117: Individuals with *F*_*ROH*_ >0.0117 (individuals who could be offspring of second cousin matings or closer matings). F>0.039: Individuals with *F*_*ROH*_ >0.039 (individuals who could be offspring of first cousin matings or closer matings, ignoring drift). F>0.093: Number and percentage of individuals with *F*_*ROH*_ >0.093 (individuals who could be offspring of avuncular matings or closer matings, ignoring drift).

| | N | Median<br>$F_{ROH}$ | IQC | F > 0.0117 | | F > 0.0391 | | F > 0.0932 | |
| --- | --- | --- | --- | --- | --- | --- | --- | --- | --- |
|  |  |  |  | N | % | N | % | N | % |
| Hunter-Gatherers | 45 | 0.0681 | 0.026 | 45 | 100 | 42 | 93.3 | 5 | 11.1 |
| West Eurasia | 43 | 0.0687 | 0.026 | 43 | 100 | 41 | 95.3 | 5 | 11.6 |
| Central Eurasia | 2 | 0.0397 | 0.006 | 2 | 100 | 1 | 50.0 | 0 | 0.0 |
| Simple Agriculturalists | 107 | 0.0287 | 0.015 | 100 | 93.5 | 18 | 16.8 | 3 | 2.8 |
| West Eurasia | 88 | 0.0291 | 0.015 | 82 | 93.2 | 14 | 15.9 | 3 | 3.4 |
| Central Eurasia | 19 | 0.0201 | 0.017 | 18 | 94.7 | 4 | 21.1 | 0 | 0.0 |
| Early Complex Agriculturalists | 237 | 0.0251 | 0.012 | 227 | 95.8 | 16 | 6.8 | 1 | 0.4 |
| West Eurasia | 151 | 0.0245 | 0.011 | 143 | 94.7 | 8 | 5.3 | 0 | 0.0 |
| Central Eurasia | 86 | 0.0267 | 0.012 | 84 | 97.7 | 8 | 9.3 | 1 | 1.2 |
| Advanced Complex Agriculturalists | 51 | 0.0136 | 0.014 | 31 | 60.8 | 2 | 3.9 | 0 | 0.0 |
| West Eurasia | 21 | 0.0012 | 0.021 | 8 | 38.1 | 1 | 4.8 | 0 | 0.0 |
| Central Eurasia | 30 | 0.0151 | 0.006 | 23 | 76.7 | 1 | 3.3 | 0 | 0.0 |
| Human Genome Diversity Panel | 448 | 0.0066 | 0.022 | 172 | 38.4 | 74 | 16.5 | 6 | 1.3 |
| West Eurasia | 139 | 0.0039 | 0.005 | 19 | 13.7 | 3 | 2.2 | 0 | 0.0 |
| Central Eurasia | 309 | 0.0156 | 0.034 | 153 | 49.5 | 71 | 23.0 | 6 | 1.9 |

We next studied the spatial distribution of *F*_*ROH*_. Notably, the distribution of *F*_*ROH*_ is highly structured in present-day Eurasia (Figure S2 B). In contrast, we found that temporal changes in *F*_*ROH*_ are largely consistent across different regions of Eurasia: neither a multiple regression analysis, with latitude and longitude as dependent variables (*β*_latitude_ = 1.9e^−04^, *p* = 0.213; *β*_longitude_ = 9.7e^−06^, *p* = 0.77), nor kriging analysis (Figure 2) revealed any prominent spatial structure for *F*_*ROH*_ through different historical categories.

### The origins of autozygosity in ancient humans

Some ancient individuals show extreme autozygosity (i.e. homozygosity created by inbreeding) within our dataset (Figure 1A). We explored the origin of these signals, asking whether, in each case, consanguinity or endogamy (i.e. genetic isolation and strong genetic drift) could be the culprit. For this, we compared the number of ROH vs. the sum of ROH per individual genome using ROH >1.5 Mb [2]. In Figure 3, the diagonal line represents an outbred population; individuals with high values along the diagonal exhibit high autozygosity due to endogamy, while “right shifts” from the diagonal indicate consanguinity. We observe that inbreeding among our sample of 440 ancient individuals can be mostly attributed to endogamy, caused by low population size. Most notably, individuals assigned to the hunter-gatherer category, with overall high *F*_*ROH*_ levels, revealed close to no indication of consanguinity. This included some West Eurasian hunter-gatherers with extreme levels of inbreeding (*F*_*ROH*_ >0.125; Chan, I0015, Villabruna, R7 and I0410). The vast majority of later-coming agriculturalists also showed no evidence of consanguinity. Within the agriculturalist sample, however, the three individuals with the most extreme levels of inbreeding (*F*_*ROH*_ >0.125) also showed clear signs of consanguinity. Based on simulations (see Materials and Methods), we estimate that NG10 from Middle Neolithic (3338-3028 cal BCE) Ireland is the offspring of an incest mating, as suggested by Cassidy and colleagues [3] (Figure 3b). We also estimate that I6671 from Early-Middle Bronze Age (3000-2039 cal BCE) Turkmenistan and I2521 from Neolithic (5619-5491 cal BCE) Bulgaria may be the offspring of avuncular matings, while also exhibiting additional autozygosity due to genetic drift.

**Figure 3.**
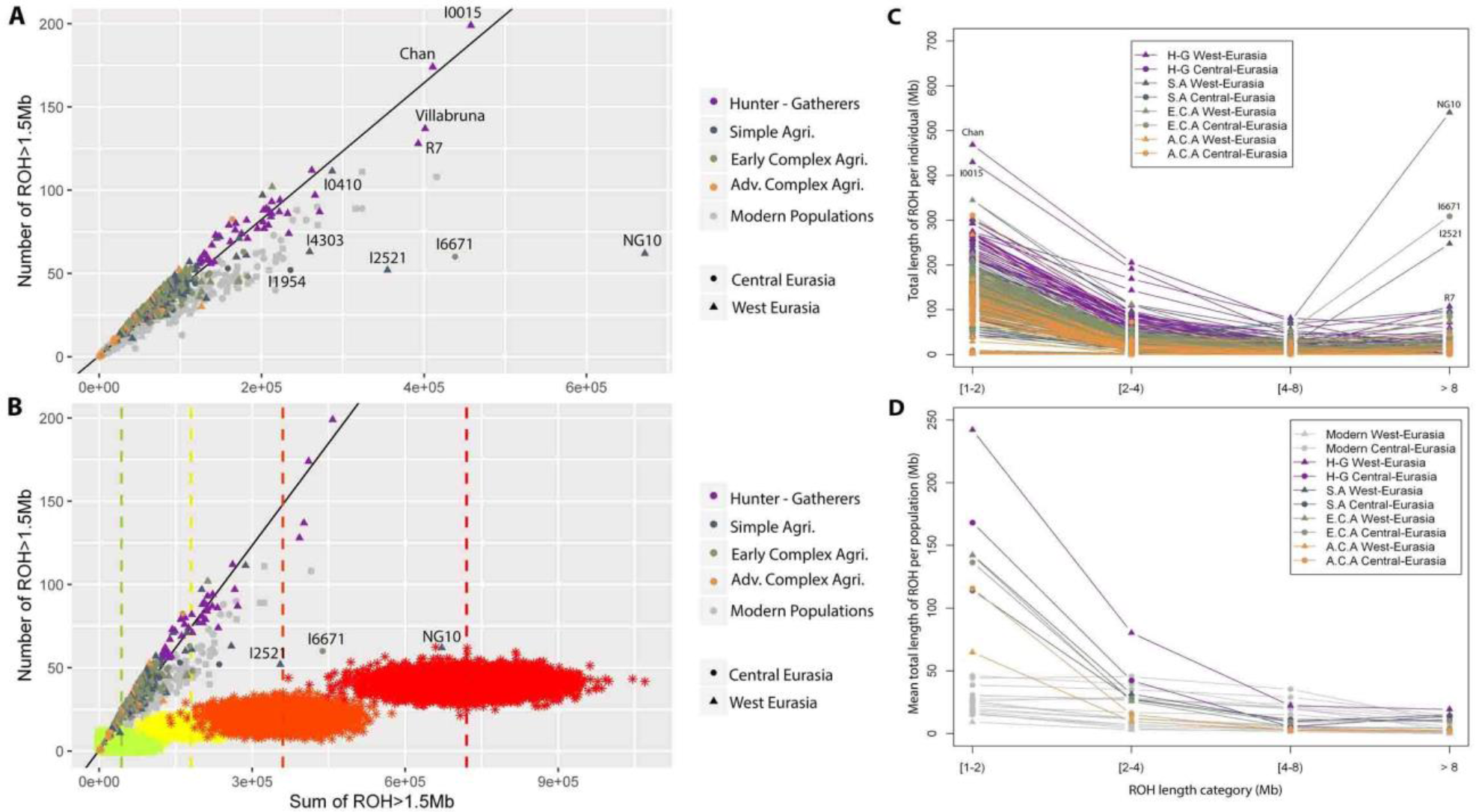
Assessing ROH origins. (**A**) Mean number of ROH and sum of ROH, for ROH >1.5 Mb, is plotted for each individual. The diagonal line is obtained by the regression of the number of ROH vs. the sum of ROH in ASW and ACB populations from the 1000 Genomes Project that represent admixed and thus outbred populations [27,28]. Consanguinity practices in the previous generation are visible as a right shift in this figure. (**B**) Simulations of the number and sum of ROH, for ROH >1.5Mb, calculated for the offspring of different consanguineous mating are shown, along with the ancient and modern samples. Asterisks points with different colours designate offspring of different consanguineous mating: second cousin (green), first cousin (yellow) avuncular (orange), incest (red). 5K simulations are represented for each consanguineous mating (see Materials and Methods). Vertical lines represent the average sum of ROH (> 1.5Mb) for the offspring of each type of consanguineous mating. **(C)** The total length of ROH (Mb) over four classes of ROH tract lengths: 1<ROH<2 Mb, 2<ROH<4 Mb, 4<ROH<8 Mb and ROH>8 Mb, described for each ancient individual. Individuals were colored according to region and period: West Eurasia hunter-gatherers (H-G West-Eurasia, shown in purple triangles), Central Eurasian hunter-gatherers (H-G Central-Eurasia shown in purple circles), West Eurasia simple agriculturalist (S.A West-Eurasia shown in blue triangles), Central Eurasian simple agriculturalist (S.A Central-Eurasia shown in blue circles), West Eurasia early complex agriculturalist (E.C.A West-Eurasia shown in green triangles), Central Eurasian early complex agriculturalist (E.C.A Central-Eurasia green circles), West Eurasia advanced complex agriculturalist (A.C.A West-Eurasia yellow triangles), Central Eurasian advanced complex agriculturalist (A.C.A Central-Eurasia yellow circles). **(D)** The total length of ROH (Mb) over four classes of ROH tract lengths as in panel C, calculated as the average for the different groups of individuals. The coloring scheme is the same as in panel C; in addition, modern-day populations are represented in grey triangles (Modern West-Eurasian populations) and circles (Modern Central-Eurasian populations).

We further studied the distribution of ROH using the total length of ROH values for different ROH track lengths (Figure 6). The size of ROH is inversely correlated with its age: longer ROH results from recent common ancestors, while shorter ROH come from distant ancestors, broken down by recombination. We found that among those hunter-gatherer individuals with extreme autozygosity created by genetic drift, total lengths of short ROH (1 Mb < ROH < 2 Mb) are high. Conversely, among the three most consanguineous individuals, NG10, I6671 and I2521, total lengths of long ROH (ROH> 8 Mb) are highest. The individual NG10 reveals 5 ROH of size >30 Mb, with an estimated age of just 1 generation [4].

Overall, we observe that consanguinity explains a small fraction of the overall autozygosity observed, with only 3 (0.6%) of the total number of the ancient individuals analysed exhibiting clear evidence of high consanguinity.

### The origins of present-day homozygosity in Central Eurasia

We then studied the spatial distribution of present-day inbreeding prevalence in relation to ancient inbreeding patterns. Figure 3D presents the average sum of the different ROH sizes across regions and historical categories. This reveals an interesting spatiotemporal structure, especially for the shorter ROH in Figure 3D. West Eurasian hunter-gatherers carry the highest total length of short ROH among all historical groups, attesting to their small population size around the early Holocene. However, this inbreeding signal is rapidly lost, and West Eurasian advanced agriculturalists carry the lowest average sum of short ROH among all ancient groups studied. In Central Eurasia, the total length of short ROH is also high in hunter-gatherers and decreases in later-coming periods, but at a more modest rate. Compared to ancient populations, present-day populations have the shortest average total length of shorter ROH, denoting large effective population size and slow genetic drift. However, this temporal pattern vanishes when we study the total length of longer ROH, e.g. ROH between 4 and 8 Mb. Importantly, ROH >4 Mb may have an age of 5 to 10 generations [5–7] and thus indicate relatively close consanguinity. Figure 3D shows that some modern Central Eurasian populations (e.g. Balochi of Pakistan or the Bedouin from Saudi Arabia), reveal higher total lengths of ROH between 4 and 8 Mb than any other group in the HGDP, as well as any of our historical categories.

In Table 1 we present comparisons of median *F*_*ROH*_ and the frequency of inbred individuals. As also observed in Figure 3, we find that contemporary populations tend to have the lowest proportions of inbred individuals. However, some contemporary Eurasian populations have exceptionally high proportions of individuals with *F*_*ROH*_ > 0.0391 (i.e. individuals who could be offspring of first cousin matings or closer matings, ignoring drift; Supplemental Information). This is especially salient among certain Central Eurasian populations. Modern groups like the Balochi, the Bedouin, or the Sindhi from Pakistan have the highest proportions of individuals with *F*_*ROH*_ >0.0391 (50%, 41.3% and 33.3% respectively).

Comparing contemporary Central vs. West Eurasia with respect to the proportion of inbred individuals, we find a significant difference between the two regions, both for individuals with *F*_*ROH*_ > 0.0391 (odds ratio = 13.5, Fisher’s exact test *p* = 9e^−10^) and also for individuals with *F*_*ROH*_ > 0.0117 (individuals who could be offspring of second cousin matings or closer matings, ignoring drift) (odds ratio = 6.2, *p* = 7e^−14^) (Table 1). Because Central Eurasian populations also exhibit relatively high total lengths of long ROH, this excess of inbred individuals could be attributed to consanguinity, rather than other processes such as caste endogamy, and is consistent with documented cultural preferences for first-cousin matings in some contemporary societies [8,9].

This raises the question whether the differential rates of consanguinity among present-day Central vs. West Eurasia could be traced back in time. In fact, we observed an excess of individuals with *F*_*ROH*_ > 0.0117 in Central vs. West Eurasia among advanced complex agriculturalists (odds ratio = 5.1, *p* = 0.009; Table 1). However, we find no indication that this was driven by consanguinity (excess of long ROH) in ancient societies (Figure 3), which suggests that the high consanguinity in this region observed today might have only a recent history.

## CONCLUSION

Our work shows that the Neolithic transition to agriculture and the emergence of complex societies did not necessarily increase the overall levels of inbreeding among humans, at least in the case of West and Central Eurasia. On the contrary, the respite from endogamy via food production and technology-driven increase in population size seem to have mitigated inbreeding levels throughout recent history. Of course here we rely on the assumption that the 440 individuals analysed in this study were representative of their time. Sampling biases caused by various factors, such as burial location of the elite vs. the commoners, or a focus on elite burials by archaeologists, could influence inferences on class-based societies. That said, as our data derive from 189 different archaeological sites and also because we observe the continuation of the same temporal trend of lower inbreeding in contemporaneous human groups, we consider our conclusions to be valid. We further note that our results are in line with previous singular reports on ROH in archaic hominins and ancient *Homo sapiens* individuals, which suggest high ROH was a common phenomenon in Paleolithic or early Holocene hunter-gatherer groups. For instance, the genome of the 50,000-year-old Altai Neanderthal individual revealed an inbreeding coefficient of ⅛, equivalent to an offspring of avuncular mating [10]. Genomes of foragers from Upper Paleolithic and Mesolithic periods from Europe and the Caucasus also display evidence for inbreeding, mainly in the form of endogamy [11,12]. Endogamy among hunter-gatherers may be expected, as it is commonly observed in wild *K*-selected species of small population size, such as mammoths [13,14].

Three points further deserve mention. One is the apparent contrast between relatively high levels of endogamic inbreeding among ancient hunter-gatherer societies, and reports of low levels of inbreeding among modern-day hunter-gatherers. Recent ethnographic studies have documented low inbreeding in a world-wide sample of contemporary hunter-gatherers living in smaller groups, compared to Amazonian horticulturalists living in larger groups [15]. Hill and colleagues also report low levels of relatedness within and high interconnection among modern-day hunter-gatherer bands [16]. This discrepancy could be attributed to various factors, such as reciprocal exogamy traditions or larger population sizes among modern-day hunter-gatherers sampled in ethnographic studies [17], which may vary from early Holocene European hunter-gatherers, which predominate our sample. In the future, estimating endogamy in non-European hunter-gatherer groups of the last 10,000 years would be crucial for resolving the prevalence of endogamy in pre-agricultural human societies. The answer, in turn, could be vital for models of how human cooperation has evolved [18,19].

Second, our data lend support, albeit weakly, to the hypothesis that extreme consanguinity may have become more common with farming. This result is also consistent with singular reports on ancient agriculturalist genomes, such as evidence for consanguinity identified in an early Neolithic farmer from Iran [20], a first-degree incest case from Neolithic Ireland [3], as well as a recent report on close-kin unions in the central Andes after 1000 CE [21]. This is intriguing as we find overall lower levels of inbreeding in agricultural than in hunter-gatherer societies, although the latter effect can be attributed to lower endogamy, and not lower consanguinity. At the same time, among the 7 most highly inbred individuals (with *F*_*ROH*_ >0.125), all 4 hunter-gatherers are mainly endogamous, while all 3 agriculturalists are mainly consanguineous. This is an unlikely observation (Fisher’s exact test *p* = 0.029) and appears consistent with the notion that consanguineous traditions could have thrived in class-based agricultural societies more readily than in more egalitarian hunter-gatherer groups [15].

Finally, we find higher consanguinity in Central vs. West Eurasia in contemporary societies. This is consistent with widespread cousin marriage practices in agricultural societies in Middle Eastern and North African (MENA) countries and in South Asia, mostly among Muslim and Jewish groups, as documented by ethnographic or genomic studies [8,22,23]. We note that cousin marriages were also common among royal dynasties and upper classes of Europe until the 20th century, and many prominent European scientists of that period are known to have married their first cousins, including Charles Darwin and Albert Einstein [24,25]. These traditions are thought to have arisen through various social factors, including the inheritance of property in class societies [15,23,26]. Interestingly, we do not observe the relatively high rates of consanguineous marriage observed in modern-day Central Eurasia in any of the past societies we studied, in Antiquity or earlier. We naturally prefer to remain cautious, especially given the limited sample size of our advanced complex agriculturalist samples from West and Central Eurasia (n=21 and n=30, respectively). Nevertheless, it appears possible that modern-day cultural patterns may have emerged late in time, possibly with the spread of Abrahamic traditions in the region.

## Acknowledgements

We are grateful to the METU CompEvo group and Torsten Günther for helpful suggestions and/or comments, Harald Ringbauer for sharing unpublished results and discussion, Jim Wilson and David Clark for support, and all archaeogenomic groups who have shared their data used in this work. This work was supported by the ERC Consolidator grant “NEOGENE” (Project No 772390 to M.S.).

## Author contributions

F.C. and M.S. conceived and designed the study; (b) F.C., K.G., and K.B.V. analysed genetic data, with contributions from D.K. and M.S.; (c) H.C.G, C.K, and Ç.A. determined the historical categories with contributions by Y.S.E.; (d) M.S. supervised the study with contributions by N.E.A., Ç.A., E.S., Y.S.E., A.Gö., F.Ö.; (e) F.C., K.G., M.S. and N.E.A. wrote the manuscript with contributions from all authors.

