## Supplementary Information for "Human inbreeding has decreased in time through the Holocene"

### Supplemental Information

### Materials & Methods

**Overview of the data, sampling and classification schemes**. We made use of the large collection of published ancient genomes in this study. We built two different datasets. The first was used for method development, and included n=44 ancient genomes with coverages higher than 10x. These were used to optimize ROH calling to suit ancient genomes with different coverage (Supplementary Dataset Table A) [1–16]. Once having determined that our approach yielded unbiased ROH estimates with genomes with coverage >3x, we built a second dataset to study the evolution of the autozygosity during the Holocene. We limited our sample to genomes of the last 15,000 years, and also to West and Central Eurasia; this spatiotemporal frame contains the highest density of published ancient genomes, providing sufficient power to test our hypotheses on change in inbreeding over time. The second dataset thus contained n=440 ancient individuals with coverages down to 3x (Supplementary Dataset Table B) [1–9,11–31]. Figure S1 shows the temporal distribution of the genomes collected, ranging between 520 CE and 12030 BCE. Further, to study the effect of sociocultural organization and economic activity, we divided ancient Eurasians into four historical categories, according to a periodization scheme described below. These included: hunter-gatherers (n=45), simple agriculturalists (n=107), early complex agriculturalist (n = 237) and advanced complex agriculturalist (n=51). For comparison purposes we also analysed 19 populations from the Human Genome Diversity Panel (HGDP) with n=444 individuals in total (Supplementary Dataset Table C). Spatial distribution of ancient and present-day individuals is shown in Figure S2. We also divided our focal region of Eurasia into two, in order to study possible spatial patterns. Following the traditional scheme, we used the Aegean, the Black Sea, the Caucasus and the Urals to delineate West and Central Eurasia. Individuals belonging to the west of this demarcation line were considered West Eurasians (n=303 ancient individuals and n=155 modern ones) and individuals belonging to the east of the line, Central Eurasians (n=137 ancient individuals and n=289 modern ones). Although alternative spatial divisions could be assumed, we note that this division overlaps with major geographical boundaries that appear to have hindered population contacts [32]. Finally, we use two outbred populations, African ancestry from Barbados in the Caribbean (ACB) and African ancestry in Southwest USA (ASW) from the 1000 Genomes dataset [33], in order to have a baseline to compare the number and sum of ROH in Figure 3.

**Processing ancient DNA.** We downloaded the published ancient human genomes from various internet sources, listed in Supplementary Dataset Tables A and B; these included data obtained by shotgun sequencing or DNA target-enrichment of 1.24 million genome-wide single-nucleotide polymorphisms (SNPs) (the so-called “1240k capture” process) [4,14,34]. To decrease bias resulting from different pipelines in the original studies, we processed all the raw data using our own pipeline. First, we mapped the reads to the human reference genome (hs37d5) using the *Burrows–Wheeler Aligner* (*BWA*, v. 0.7.15) [35], with parameters '-l 16500, -n 0.01, -o 2'. Then we used *SAMtools* "*merge*" (v. 1.9) [36] to merge all libraries from the same individual and removed PCR duplicates using *FilterUniqueSAMCons.py* [37]. We removed reads shorter than 35 base pairs and those with >10% mismatches to the reference genome. We trimmed the BAM files of the ancient individuals to remove the postmortem damage at the ends to avoid interpreting them as true variants. We trimmed reads from both ends by 10 bp if no UDG treatment was used in the original study, and 2 bp if the samples were UDG-treated, using the *trimBAM* command of *bamUtil* software [38]. We used *SAMtools mpileup* to call genotypes of ancient individuals using the 1240k SNP panel as reference (parameters: '-B -q30 -Q30') [4]. The output BCF files were converted to VCF files with parameters '-mV indels' using *bcftools call* command [39]. We obtained the mean coverage per genome across the 1240K SNP set by collecting SNP read depths from each VCF file (i.e. the 'DP' field). Finally, the VCF files were converted to *PLINK* input files using *PLINK v1.9* [40,41].

**Processing modern data.** In an attempt to enrich the dataset, we merged the above ancient individuals with 444 individuals from the Human Genome Diversity Panel (HGDP) (belonging to 19 populations) [42] and 235 admixed (outbred) individuals from the 1000 Genomes (belonging to two populations: ASW and ACB) [33]. We used the version 3.0 of the HGDP which consists of high coverage WGS (data obtained from ftp://ngs.sanger.ac.uk:21/production/hgdp/hgdp_wgs.20190516) and genotype data (Illumina BedStation and Infinium Omni 2.5) from the 1K Genomes individuals (data obtained from http://ftp.1000genomes.ebi.ac.uk/vol1/ftp/relea se/20130 502). In order to have a homogeneous dataset we filtered both HGDP and 1K Genomes individuals using the 1240k SNP panel. Thus, all the individuals used in this study had the same SNP panel. Since HGDP data is based on the GRCh38 reference genome, but 1240K is based on GRCh37, a liftover to the HGDP dataset was applied.

**Statistical analyses.** All statistical tests and procedures described below were performed using *R v3.6.1* [43]. All tests were conducted two-sided.

**ROH calling procedures.** We used *PLINK v1.9* [40,41] to identify ROH. *PLINK* has been extensively used to call ROH in human genomic studies and it is the most widely used software [44]. We decided to use *PLINK* due to its methodological advantages and for comparing its performance with other software utilized in previously published studies. Its direct observational approach allows the researcher to have full control of the ROH calling process. Also by modifying its parameters, it has been shown that it is possible to obtain equivalent ROH estimations between different sequence technologies and genomic coverages [44]. ROH were called in the autosomal genome.

The following conditions where used:

*--homozyg-snp 50.* Minimum number of SNPs that a ROH is required to contain (50 SNPs).

*--homozyg-kb 300.* Length in Kb of the sliding window (300 Kb).

*--hmozyg-density 50.* Required minimum density to consider a ROH (1 SNP in 50 Kb).

*--homozyg-gap 1000.* Length in Kb between two SNPs to be considered in two different segments (1 Mb).

*--homozyg-window-snp 50.* Number of SNPs that the sliding window must have (50 SNPs).

*--homozyg-window-het (0 - 1).* Number of heterozygous SNPs allowed in a window (0 or 1).

*--homozyg-window-missing 5.* Number of missing calls allowed in a window (5 calls).

*--homozyg-window-threshold 0.05.* Proportion of overlapping windows that must be called homozygous to define a given SNP as in a "homozygous" segment (5%).

We collected the following statistics from the outcome file of each *PLINK* run:

(a) the total number of ROH events for ROH longer than 1 Mb ($NROH_{>1Mb}$),

(b) the total number of ROH events for ROH shorter than 1 Mb ($NROH_{<1Mb}$),

(c) the total sum of ROH lengths for ROH longer than 1 Mb ($SROH_{>1Mb}$),

(d) the total sum of ROH lengths for ROH shorter than 1 Mb ($SROH_{<1Mb}$),

(e) the total sum of ROH longer than 1.5 Mb divided by the total length of the autosomal genome, defined as genomic inbreeding coefficient, or $F_{ROH}$ (McQuillan et al. 2008).

On each genome we estimated ROH using *PLINK* in two ways, allowing 0 heterozygous SNP per window ('het 0'), and allowing 1 heterozygous SNP ('het 1'). In addition, for each of the statistics above, we calculated the average between the 'het 0' estimate and the 'het 1' estimate, and we refer to this average ROH estimate as 'het 0.5' (Figure S3). For instance, the $SROH_{>1Mb}$ value for 'het 0.5' is the average between $SROH_{>1Mb}$ for 'het 0' and $SROH_{>1Mb}$ for 'het 1'.

**Studying the effects of varying coverage on variant and ROH calling by simulation**. To study the effect of genome coverage in variant calling and downstream ROH estimation, we simulated low coverage genomes by downsampling the n=44 ancient genomes with >10x coverage (Supplementary Dataset Table A) into 10x, 5x, 3x and 2x genomes, yielding a total of n=220 full or partial genomes. For this, we used *Picard*'s *DownsampleSam* tool (https://broadinstitute.github.io/picard/) to randomly extract aligned reads from the BAM files to obtain 10x, 5x, 3x and 2x coverage versions. Table S1 shows variant calling results across coverages. It is possible to see that downsampling increases the frequency of homozygous SNPs.

Consequently, downsampling leads to systematic overestimation of ROHs, as can be seen in Figure S3.

**An empirical approach to remove the effect of coverage variance on ROH calls**. To assess the effects of different genomic coverage and the number of heterozygous SNPs allowed per window on the sum and number of ROH, we called ROH by alternative approaches, and studied the effects using regression analyses. First, using the standard approach, we calculated ROH using the 'het 0', 'het 1' and 'het 0.5' schemes on all genomes. We then tested the effects of coverage and ROH calling scheme by fitting the below general linear model, or model 1:

$Y_{ijk} = \gamma_{00} + C_i + V_j + U_k$,

where $Y_{ijk}$ is the variable response (number or sum of ROH, longer or shorter than 1 Mb) with coverage $i$, the number heterozygous SNPs allowed per window $j$, and the individual (as random effect) $k$. $\gamma_{00}$ is the overall average media, $C_i$ is the fixed effect of the genomic coverage of each individual, $V_j$ is the fixed effect of the number of heterozygous SNPs allowed per window and $U_k$ is the random effect of the individuals.

In the second, empirical approach, we fixed the number of heterozygous SNPs allowed per window depending on coverage, as follows: (a) for genomes with average genomic coverage ≥4x we allowed 1 heterozygous SNP per window (i.e. calculated ROH using the 'het 1' scheme above), (b) for genomes with average genomic coverage <4x we used the 'het 0.5' scheme above. In other words, we corrected for the observed effect of excess homozygosity caused by lower coverage, by calculating ROH estimates as averages between 'het 0' and 'het 1'. We then fit the following general linear model 2:

$Y_{ijk} = \gamma_{00} + C_i + U_j$,

where $Y_{ijk}$ is the variable response (number and sum of ROH longer and shorter than 1 Mb) with coverage $i$, and individual (as random effect) $j$. $\gamma_{00}$ is the overall average media, $C_i$ is the fixed effect of the genomic coverage of each individual, and $U_j$ is the random effect of the individual.

The results of both models 1 and 2 are shown in Tables S1 and S3. We found that coverage was highly significant ($P$<2.2e-16) for both models. However, when the different coverage classes (>10x, 10x, 5x, 3x and 2x) are pairwise compared (Table S4, Figure S2) we can see that, by using the empirical approach, it is possible to obtain similar (statistically non-different) estimations of the sum and number of ROH (for ROH >1Mb) for average genomic coverage down to 3x.

To confirm that genomic coverage does not influence ROH calls using this empirical, we called ROH >1Mb across 440 ancient individuals with genome coverages ≥3x (see above), using the empirical method. We then calculated Kendall's rank correlation between genomic coverage vs. $NROH_{>1Mb}$, $SROH_{>1Mb}$ and average time (BCE). Neither correlation was significant ($z$ = -1.61, $p$ = 0.107, $z$ = -1.72, $p$ = 0.088 and $z$ = -1.67, $p$ = 0.094, respectively). However, for ROH <1Mb, none of these approaches could remove the effect of variable genomic coverage on ROH calls, i.e. $NROH_{<1Mb}$ and $SROH_{<1Mb}$ statistics.

**Comparison with alternative approaches for ROH calls on low coverage genomes.** We compared our results with those of Ringbauer et al. [45], who used a fundamentally different approach to detect the ROH, in which they modeled pseudo-haploidized genotypes of ancient individuals as a mosaic fitting a reference panel of multiple modern haplotypes using linkage disequilibrium information and employing a Hidden Markov Model. We found that the sum of ROH and number of ROH values for the n=384 samples used in both studies and four different categories (ROH > 4 cM; ROH > 8 cM; ROH >12 cM; ROH > 20 cM) are highly similar (data not shown). The consistency between our results, despite the difference in approaches, increases the reliability of both studies.

***$F_{ROH}$* measurements.** In view of our downsampling experiments and the regression analysis results, in downstream analyses we used the genomic inbreeding coefficient, or $F_{ROH}$ (sum of ROH >1.5 Mb) (McQuillan et al. 2008), calculated using our empirical approach: i.e. using the 'het 1' scheme for genomes <4x coverage, and calculated using the 'het 0.5' scheme for genomes between 3x to 4x coverage. Genomes with <3x coverage were not used. We used different $F_{ROH}$ thresholds to define different consanguinity matings (Table 1). $F_{ROH}$ > 0.0117 (average value between the genealogical inbreeding coefficient of second and third cousin) designates individuals who could be the offspring of a second cousin marriage. $F_{ROH}$ > 0.039 (average value between the genealogical inbreeding coefficient of first and second cousin) designates individuals who could be the offspring of a first cousin marriage. $F_{ROH}$ >0.093 (average value between the genealogical inbreeding coefficient of first and avuncular mating) designates individuals who could be the offspring of an avuncular marriage.

**Temporal and spatial distribution.** We used different approaches to assess the effect of temporal and spatial distribution in $F_{ROH}$ of the individuals analysed in this study. We first fit a simple regression analysis of $F_{ROH}$ on the archaeological age of ancient individuals. We also compared among different historical categories using a Wilcoxon pairwise rank-sum test with continuity correction using the R 'wilcox.test' function. We further fit a multiple regression model with both the archaeological age and historical category. To test the spatial distribution, we first fit a multiple regression analysis with $F_{ROH}$ as dependent variable, and the longitude, latitude, and archaeological age of each ancient individual as independent variables. To delve deeper in the spatial distribution of the $F_{ROH}$, we divided the complete dataset into the designed historical categories, and we fitted a kriging, or Gaussian process regression, using the R functions *variogram()*, *fit.variogram()* and *krige()* in package *Gstat v2.06* [46]. To obtain the variogram we used an exponential model.

**Origins of autozygosity.** In order to assess the origins of the autozygosity exhibited by the ancient individuals we used the number and sum of ROH longer than 1.5Mb, as explained by Ceballos and colleagues [47]. However, we went a step forward by simulating the number and sum of ROH of different consanguineous mating. We simulated 20K second cousins, first cousins, avuncular (uncle niece or double first cousin) and incest mating (5K each) by using the GRCh37 genetic map and an inhouse R script.

**Historical periodization based on social complexity.** Identifying social complexity in archaeology is an issue in itself, and the debate in archaeology and cultural anthropology still continues [48–51]. Social complexity can be defined as increased differences in status that leads to social hierarchies in control of economic and social activities by a centralized agent and its bureaucratic devices, such as a chief or a state [52]. Archaeological correlates to this phenomenon include increased population size as demonstrated in settlement patterns, complex organization of labor as demonstrated in labor intensive subsistence activities and architectural projects, social networks as demonstrated in long-distance exchange of artifacts and raw materials, or intensified and elaborated ritual activities [53]. Here, we use a fourfold division that corresponds to some of the most important socio-economic thresholds in human (pre)history.

Hunter-gatherer groups are small-scale egalitarian mobile bands that subsist by utilizing the wild resources in the land. While some hunter-gatherer communities can be involved in long-distance exchange, elaborate ritual, and labor control for large scale architectural projects [54], the emergent complexity is often fragile in the absence of storage and redistributive mechanisms [55]. In these groups, often the accumulation of wealth by individuals or certain groups as well as transference of acquired status to kin members are not permitted.

Prior to the Holocene, hunting-gathering-foraging was the sole source of livelihood; however, with the onset of the Holocene, long term sedentism and intensification of animal and plant management gave way to domestication and the establishment of agricultural societies, i.e. the Neolithic Process [56]. It is understood that the advent of agriculture is the basis for more solid forms of social complexity, particularly due to the emergence of a new social and economic organization that this pattern of subsistence requires. In particular, the increase in population sizes, the emergence of storage and redistribution mechanisms, along with a new social and economic organization that is focused on competing households and extended kinship networks, together seem to have resulted in more complex forms of social organization, such as chiefdoms and states [57–59]. In these societies, competing groups seek to expand their labor force within a kinship making system, which can be secured by marriage alliances with a wide range of groups; in this context, prestigious families attract more alliances and multiple spouses can be attained.

For our purposes, we identified three types of agriculturalist societies. We chose to name the early sedentary agricultural villages without a centralized institution that is at the top of a hierarchical social organization as simple agriculturalists, in any geographical area [60,61]. These societies represent the earliest farming groups in their region, either growing organically from the preceding hunter-gatherer groups, or through a process of drawn-out interaction and/or replacement by pre-existing farmer communities. Secondly, the groups that show forms of incipient centralization and institutionalized hierarchy are named early complex agriculturalists, indicating a degree of social complexity as particularly evidenced in their ritual activity, settlement pattern, architecture and craft specialization [62–66]. These include those societies caught in the process of urbanization, early metal-working societies, and pastoral groups that eventually boosted their mobility with the domestication of the horse. Lastly, the advanced complex agriculturalists include the highly stratified societies organized around states, depending on a system of economical accumulation and redistribution based on a formal, typically hereditary leadership [67–69].

Table S5 provides a description of the rough initiation dates of each historical category in each of the two regions of Eurasia.

### Supplemental Tables

**Table S1.** Variant calling comparison of different average genomic coverage among 44 ancient individuals.

| Cov. | % of SNP | Concor. % | Discor. % | Het -> Hom | Hom -> Het |
|---|---|---|---|---|---|
| 10x | 97.44 | 95.53 | 4.47 | 88.30 | 11.70 |
| 5x | 91.56 | 90.47 | 9.53 | 92.25 | 7.75 |
| 3x | 87.69 | 87.02 | 12.98 | 93.77 | 6.23 |
| 2x | 75.46 | 80.23 | 20.77 | 98.64 | 1.36 |

We downsampled 44 ancient individuals (Supplementary Dataset Table A) with an average genomic coverage >10x to obtain 10x, 5x, 3x and 2x versions, shown in column “Cov”. “% of SNP”: average percentage of SNPs called in downsampled versions relative to the original individual. The variant calling concordance and discordance of the results obtained for each genomic coverage with respect to the original genotype, in percentage, is shown in columns titled “Concor. %” and “Discor. %”, respectively. Among the discordant sites the percentage of those that in the original where heterozygous but in the downsampled version turned homozygous, and vice versa, are shown in columns titled “Het -> Hom” and “Hom -> Het”, respectively.

**Table S2:** Results of pairwise Wilcoxon rank-sum test with continuous correction of the genomic inbreeding coefficient (FROH) among the five historical categories. P-values are not corrected for multiple testing.

| | **W** | **P-value** |
|---|---|---|
| Hunter-Gatherers vs Simple agriculturalists | 4369 | 2.48e-15 |
| Hunter-Gatherers vs Early complex agriculturalists | 10209 | < 2.2e-16 |
| Hunter-Gatherers vs Adv. complex agriculturalists | 2258 | 3.63e-16 |
| Hunter-Gatherers vs Modern-day populations | 18675 | < 2.2e-16 |
| Simple agriculturalists vs Early complex agriculturalists | 14105 | 0.09514 |
| Simple agriculturalists vs Adv. complex agriculturalists | 4469.5 | 9.6e-11 |
| Simple agriculturalists vs Modern-day populations | 36376 | < 2.2e-16 |
| Early complex agriculturalists vs Adv. complex agriculturalists | 9894 | 9.6e-13 |
| Early complex agriculturalists vs Modern-day populations | 79083 | < 2.2e-16 |
| Adv. complex agriculturalists vs Modern-day populations | 12226 | 0.4117 |

**Table S3.** Correction scheme to remove the effect of coverage variance on ROH calls.

| | Sum of ROH (ROH>1Mb) | | Number of ROH (ROH>1Mb) | |
|---|---|---|---|---|
| **Model 1 Effects** | **F-Value** | **P-value** | **F-Value** | **P-value** |
| **Coverage (fixed)** | 100.78 | <2.2E-16 | 118.02 | <2.2E-16 |
| **HET (fixed)** | 292.2 | <2.2E-16 | 230.85 | <2.2E-16 |
| **Model 1 Random Effects** | **LogLik** | **P-value** | **LogLik** | **P-value** |
| **Individual (random)** | -8409.1 | <2.2E-16 | -327.2 | <2.2E-16 |
| **Model 2 Fixed Effects** | **F-Value** | **P-value** | **F-Value** | **P-value** |
| **Coverage** | 41.79 | <2.2E-16 | 60.01 | <2.2E-16 |
| **Model 2 Random Effects** | **LogLik** | **P-value** | **LogLik** | **P-value** |
| **Individual** | -2277.3 | <2.2E-16 | -1073.5 | <2.2E-16 |

Outcomes are shown for sum and number of ROH >1 Mb. Model 1 includes ROH calls made using 0 and 1 heterozygous SNP allowed per window ('het 0' and 'het 1), along with the average between those ('het 0.5'). "Coverage" is a quantitative variable describing the mean SNP coverage per genome, "HET" is a qualitative variable describing the heterozygosity scheme used. "Individual" is included in the models as a random effect. In model 2 the heterozygous SNP allowed per window is fixed (and therefore "HET" is not included"). ROH calls are made using 'het 1' for coverages >4x and 'het 0.5' for coverages between 4x-2x. "LogLik": log likelihood.

**Table S4.** Model 1 and Model 2 pairwise comparison through the least squares means of the different genomic coverages. Non-significant results (i.e. cases where coverage does not have an effect on sum or number of ROH) are highlighted in grey.

| | **Sum of ROH (ROH>1Mb)** | | | **Number of ROH (ROH>1Mb)** | | |
|---|---|---|---|---|---|---|
| **Model 1** | **Estimate** | **Std.Error** | **p-value** | **Estimate** | **Std.Error** | **p-value** |
| **>10x-10x** | -11272.9 | 9968.4 | 0.25857 | -3.84884 | 4.21839 | 0.3619 |
| **>10x-5x** | -6942.2 | 9968.4 | 0.48644 | -4.18605 | 4.21839 | 0.3214 |
| **>10x-3x** | -47469.5 | 9968.4 | 2.40E-06 | -22.90698 | 4.21839 | 8.20E-08 |
| **>10x-2x** | -161053.3 | 9968.4 | <2.2E-16 | -74.53488 | 4.21839 | <2.2E-16 |
| **10x-5x** | -18215.1 | 9968.4 | 0.06816 | -8.03488 | 4.21839 | 0.0573 |
| **10x-3x** | -58742.4 | 9968.4 | 6.35E-09 | -26.75581 | 4.21839 | 4.46E-10 |
| **10x-2x** | -172326.2 | 9968.4 | <2.2E-16 | -78.38372 | 4.21839 | <2.2E-16 |
| **5x-2x** | -40527.3 | 9968.4 | 5.43E-05 | 18.72093 | 4.21839 | 1.08E-05 |
| **5x-3x** | -154111.1 | 9968.4 | <2.2E-16 | 70.34884 | 4.21839 | <2.2E-16 |
| **3x-2x** | 113583.8 | 9968.4 | <2.2E-16 | 51.62791 | 4.21839 | <2.2E-16 |
| **Model 2** | **Estimate** | **Std.Error** | **p-value** | **Estimate** | **Std.Error** | **p-value** |
| **>10x-10x** | -6194.8 | 12055.2 | 0.60802 | -2.046512 | 5.019349 | 0.683995 |
| **>10x-5x** | -16509.1 | 12055.2 | 0.17268 | -7.883721 | 5.019349 | 0.118142 |
| **>10x-3x** | -13809.6 | 12055.2 | 0.25362 | -13.05814 | 5.019349 | 0.071 |
| **>10x-2x** | -127393.3 | 12055.2 | <2.2E-16 | -64.68605 | 5.019349 | <2.2E-16 |
| **10x-5x** | -22703.9 | 12055.2 | 0.121 | -9.930233 | 5.019349 | 0.09 |
| **10x-3x** | -20004.4 | 12055.2 | 0.235 | -15.10465 | 5.019349 | 0.09 |
| **10x-2x** | -133588.1 | 12055.2 | <2.2E-16 | -66.73256 | 5.019349 | <2.2E-16 |
| **5x-3x** | -2699.6 | 12055.2 | 0.82308 | 5.174419 | 5.019349 | 0.304072 |
| **5x-2x** | -110884.2 | 12055.2 | <2.2E-16 | 56.802326 | 5.019349 | <2.2E-16 |
| **3x-2x** | -113583.8 | 12055.2 | <2.2E-16 | 51.627907 | 5.019349 | <2.2E-16 |

**Table S5:** Approximate initiation dates of different social complexity categories for different regions [70–72].

| Regions | Simple Agriculturalists | Early Complex Agriculturalists | Advanced Complex Agriculturalists |
|---|---|---|---|
| Near East | 10$^{th}$-8$^{th}$ mil. BC | 5$^{th}$-4$^{th}$ mil. BC | 3$^{rd}$-2$^{nd}$ mil. BC |
| SE Europe | 7$^{th}$ mil. BC | 4$^{th}$ mil. BC | 2$^{nd}$ mil. BC |
| Central Asia | 7$^{th}$ mil. BC | 4$^{th}$ mil. BC | 2$^{nd}$-1$^{st}$ mil. BC |
| Central Europe | 6$^{th}$ mil. BC | 3$^{rd}$ mil. BC | 1$^{st}$ mil. BC |
| West Mediterranean | 6$^{th}$ mil. BC | 3$^{rd}$ mil. BC | 1$^{st}$ mil. BC |
| East Europe | 6$^{th}$-5$^{th}$ mil. BC | 4$^{th}$ mil. BC | 1$^{st}$ mil. BC |
| Atlantic Europe | 5$^{th}$ mil. BC | 3$^{rd}$ mil. BC | 1$^{st}$ mil. BC |
| British Isles | 4$^{th}$ mil. BC | 3$^{rd}$ mil. BC | 1$^{st}$ mil. BC |
| North Europe | 4$^{th}$-3$^{rd}$ mil. BC | 3$^{rd}$ mil. BC | 1$^{st}$ mil. BC |
| Siberia | - | 4$^{th}$ mil. BC | 1$^{st}$ mil. BC |

**Supplementary Figures**

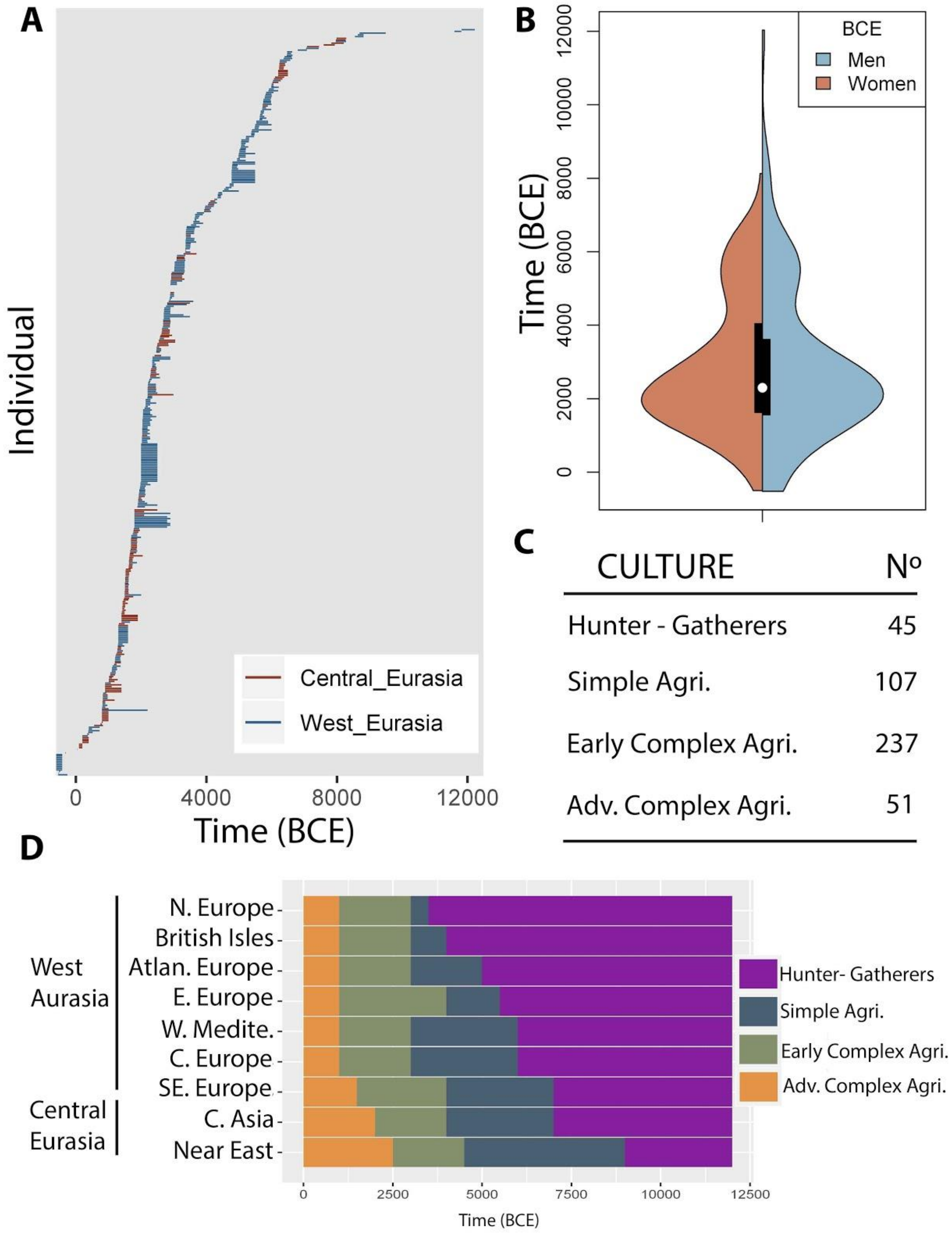

**Figure S1.** Temporal distribution of the ancient individuals analysed. **A)** Upper and lower BCE estimations for the different individuals. Colours design two geographical regions, West Eurasia and Central Eurasia divided by the Carpathian and Urals Mountains (see Materials and Methods). **B)** Violin plot of the average estimated BCE for ancient men and women. **C)** Ancient individuals were grouped into four historical categories: Hunter-gatherers, simple agriculturalists (Simple

Agri.), early complex agriculturalists (Early Complex Agri.), and advanced complex agriculturalists (Adv. Complex Agri.) (see Materials and Methods). **D)** Time scale of the four historical categories in different regions of West and Central Eurasia.

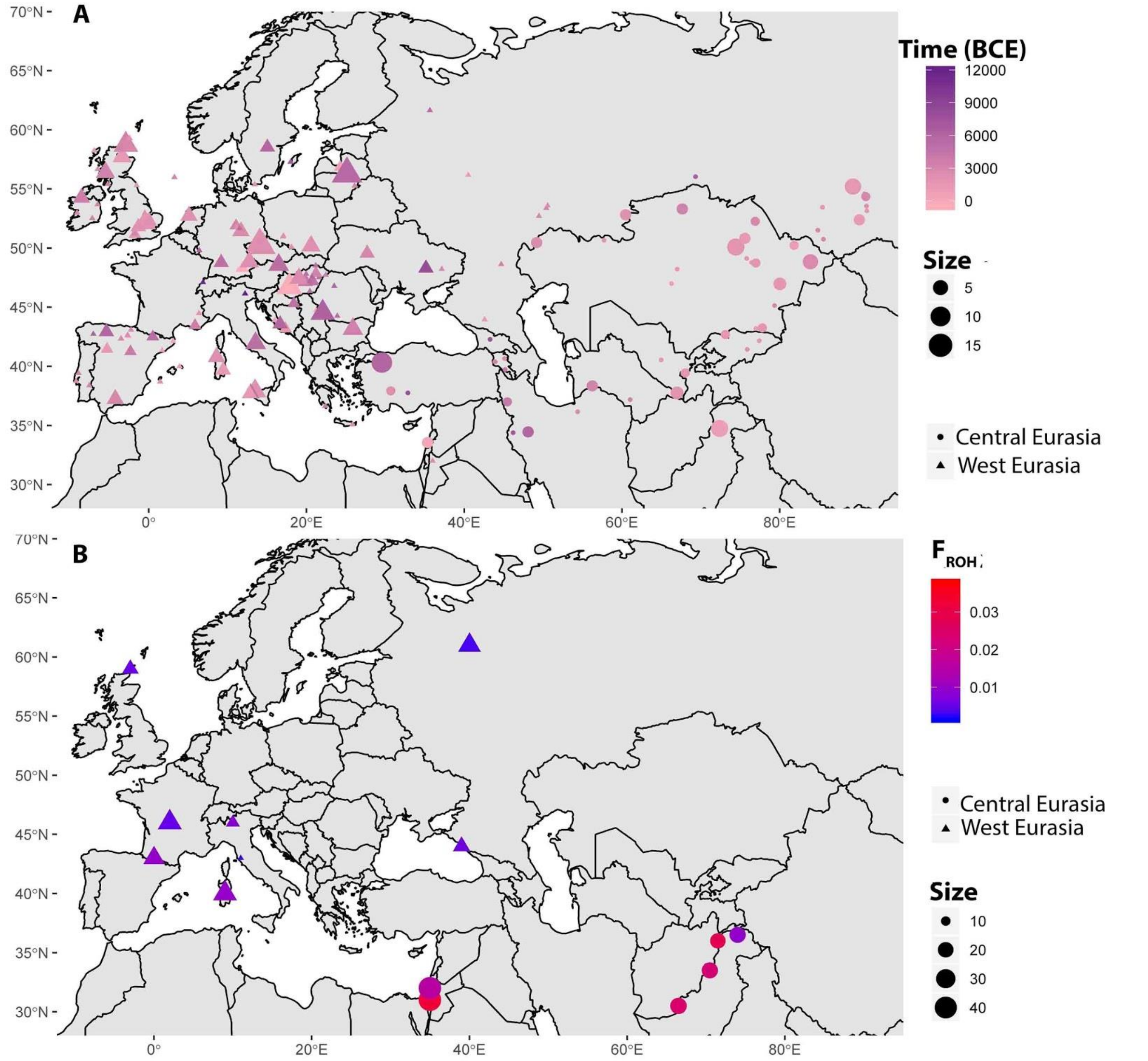

**Figure S2. Spatial distribution of ancient and modern individuals.** Populations and individuals were grouped into two classes according to their geographic location: West (triangles) and Central (circles) Eurasia (see Materials and Methods)**. A)** Spatial distribution of ancient samples. Colour represents the average BCE of the archaeological site. Size is proportional to the number of individuals sampled in each archaeological site. **B)** Spatial distribution of modern individuals from the HGDP. Colour represents the population's average genomic inbreeding coefficient ($F_{ROH}$). Size designates the sample size of the population.

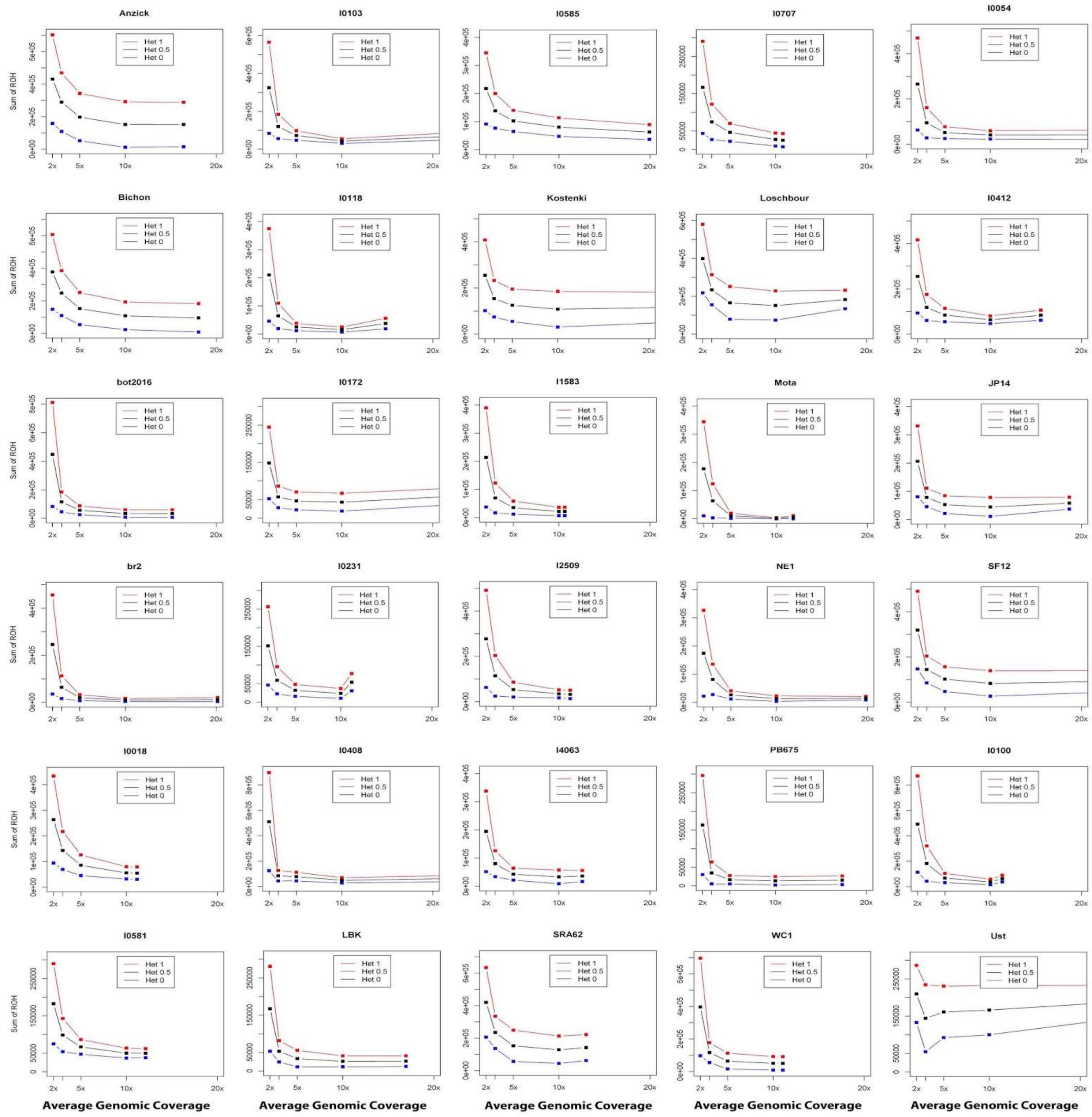

**Figure S3. Sum of ROH for different mean genomic coverage in 35 ancient individuals.** The total sums of ROH (>1 Mb) of 35 individuals with an average genomic coverage larger than 10x, are represented along with 10x, 5x, 3x and 2x downsampled versions. The plot also shows the sums of ROH for different heterozygous SNPs allowed per window. ROH estimates were obtained allowing either 1 heterozygous SNP per window ('het 1', red lines) or 0 heterozygous SNP per window ('het 0', blue lines), and also by calculating the average between 'het 0' and 'het 1' ROH estimates ('het 0.5', black lines).

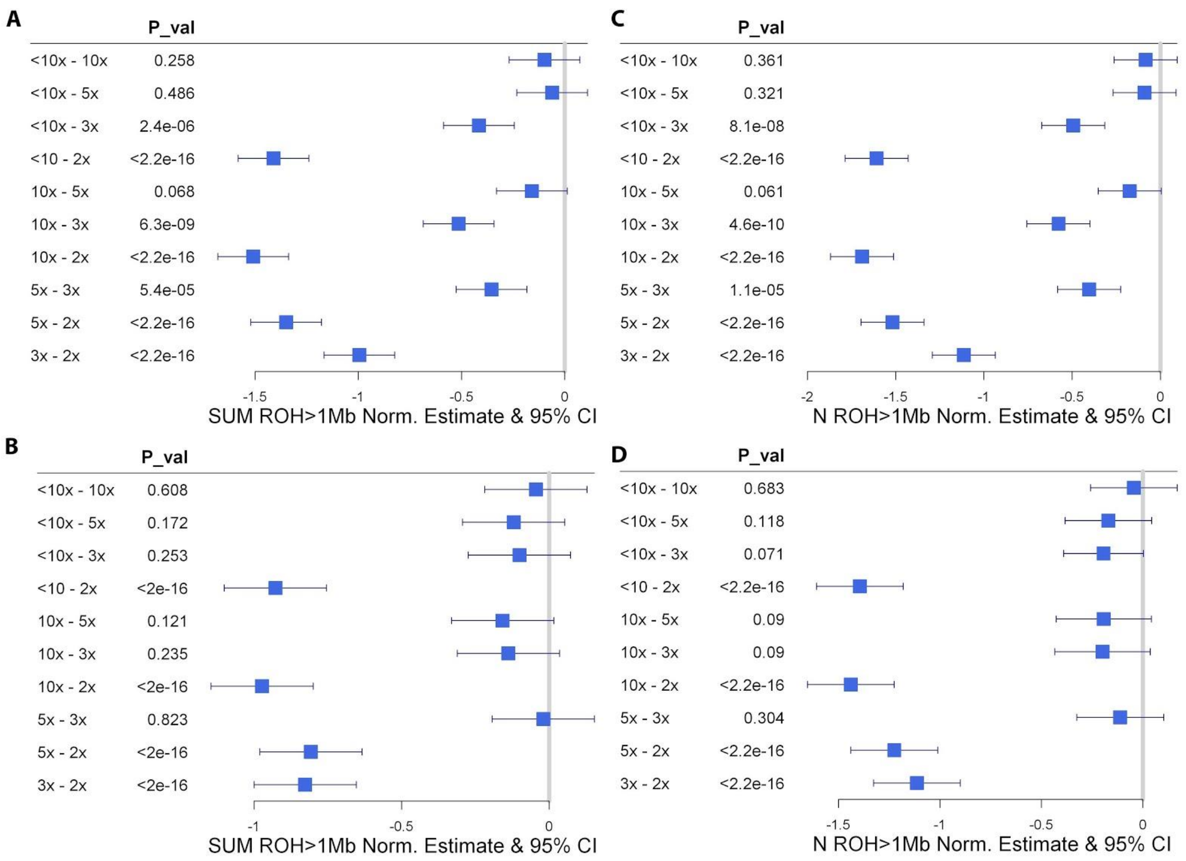

**Figure S4. Effect of the genomic coverage on the sum and number of ROH longer than 1 Mb.** To assess the effect of the coverage on the estimation of the number and sum of ROH (>1 Mb) we used different *PLINK* conditions. In this figure, we represent the effect of the pairwise change between two different genomic coverages: coverage larger than 10x and 10x (>10x – 10x), larger than 10x and 5x (>10x – 5x), larger than 10x and 3x (>10x – 3x), larger than 10x and 2x (>10x – 2x), 10x and 5x (10x – 5x), 10x and 3x (10x – 3x), 10x and 2x (10x – 2x), 5x and 3x (5x – 3x), 5x and 2x (5x – 2x), and finally, 3x and 2x (3x – 2x). To allow comparison between pairs of coverages effect estimates and the 95% confidence intervals are shown in units of within-group standard deviations. **A)** Effect of the coverage on the sum of ROH >1 Mb, considering a model where 0 and 1 heterozygotes SNP are allowed for each individual analysed as also the average of the sum of ROHs obtained allowing 0 heterozygous SNP per window or 1 heterozygous SNP per window ('het 0.5') (see Materials and Methods). **B)** Effect of the coverage on the sum of ROH >1 Mb considering a model with fixed heterozygotes SNP per window. The model used the sum of ROH obtained allowing 1 heterozygous SNP per window for individuals with a genomic coverage above 4x ('het 1'), and the average of the sum of ROHs obtained allowing 0 heterozygous SNP per window or 1 heterozygous SNP per window ('het 0.5'), for individuals with a genomic coverage below 4x (see Materials and Methods). **C)** Effect of the coverage on the number of ROH >1 Mb considering a model where 0, 1 and 0.5 ('het 0.5') heterozygotes SNP are allowed for each individual analysed. **D)** Effect of the coverage on the number of ROH longer than 1 Mb considering a model with fixed heterozygotes SNP per window. The model used the sum of ROH obtained allowing 1 heterozygous SNP per window for individuals with a genomic coverage above 4x ('het 1'), and the average of the sum of ROHs obtained allowing 0 heterozygous SNP per window or 1 heterozygous SNP per window ('het 0.5'), for individuals with a genomic coverage below 4x (see Materials and Methods).

**Supplemental Dataset**

**Supplemental Dataset Table A: List of genomes used in downsampling simulations.** We used 44 individuals' published genomes with >10x average genomic coverage to study the effects of varying coverage on variant and ROH calling by randomly downsampling genomic data. The table shows individual IDs according to the relevant publication (*IID*), and also the average coverage at 1240K SNPs and coverage standard deviation of each genome (*Average coverage* and *Coverage SD*, respectively). The table also shows the different probabilities used to obtain *10x*, *5x*, *3x* and *2x* genomic coverage versions of each individual's genome, using the Picard software DownsampleSam tool, which retains a random subset of reads according to a given probability (columns D to G). The table also presents information collected from the relevant publications: the upper and lower limits of published calibrated radiocarbon age (*DP1* and *DP2* respectively), the country of origin (*Country*), the burial site (*Site*), the latitude and longitude (*Lat* and *Long*) of each site, and finally, the paper that released the genomic information of each individual (*Publication*). Radiocarbon age is given in years BCE, with negative values indicating years CE.

**Supplemental Dataset Table B: List of ancient genomes and their FROH estimates.** The table lists individual IDs according to the relevant publication (*IID*) of the 440 ancient individuals with >3x average genomic coverage used in the study. The table lists the historical category each individual was assigned (*Historical category*), as well as information collected from the relevant publications: the upper and lower limits of published calibrated radiocarbon age (*DP1* and *DP2* respectively), the average calibrated radiocarbon age [*Age (BCE)*], the region, country and burial site of origin (*Region, Country, Site*) the latitude and longitude of each burial site (*Lat* and *Long*), the sex of the individual (*Sex*), the average genomic coverage at 1240K SNPs (*Average coverage*), the calculated genomic inbreeding coefficient ($F_{ROH}$), and the publication that released the genomic information of each individual (*Publication*). Radiocarbon age is given in years BCE, with negative values indicating years CE.

**Supplemental Dataset Table C: List of modern genomes and their FROH estimates.** The table lists individual IDs (*IID*) of 448 modern individuals from the HGDP dataset used in this study. The table shows the population of origin (*Pop*), the latitude and longitude of the sampling site (*Lat* and *Long*, respectively), the region of origin (*Region*) and the sex of each individual (*Sex*), as well as the calculated genomic inbreeding coefficient ($F_{ROH}$).
